## Supplementary figures and images for "Smith-Magenis syndrome protein RAI1 regulates body weight homeostasis through hypothalamic BDNF-producing neurons and neurotrophin downstream signalling"

### Figure 1 suppl 1

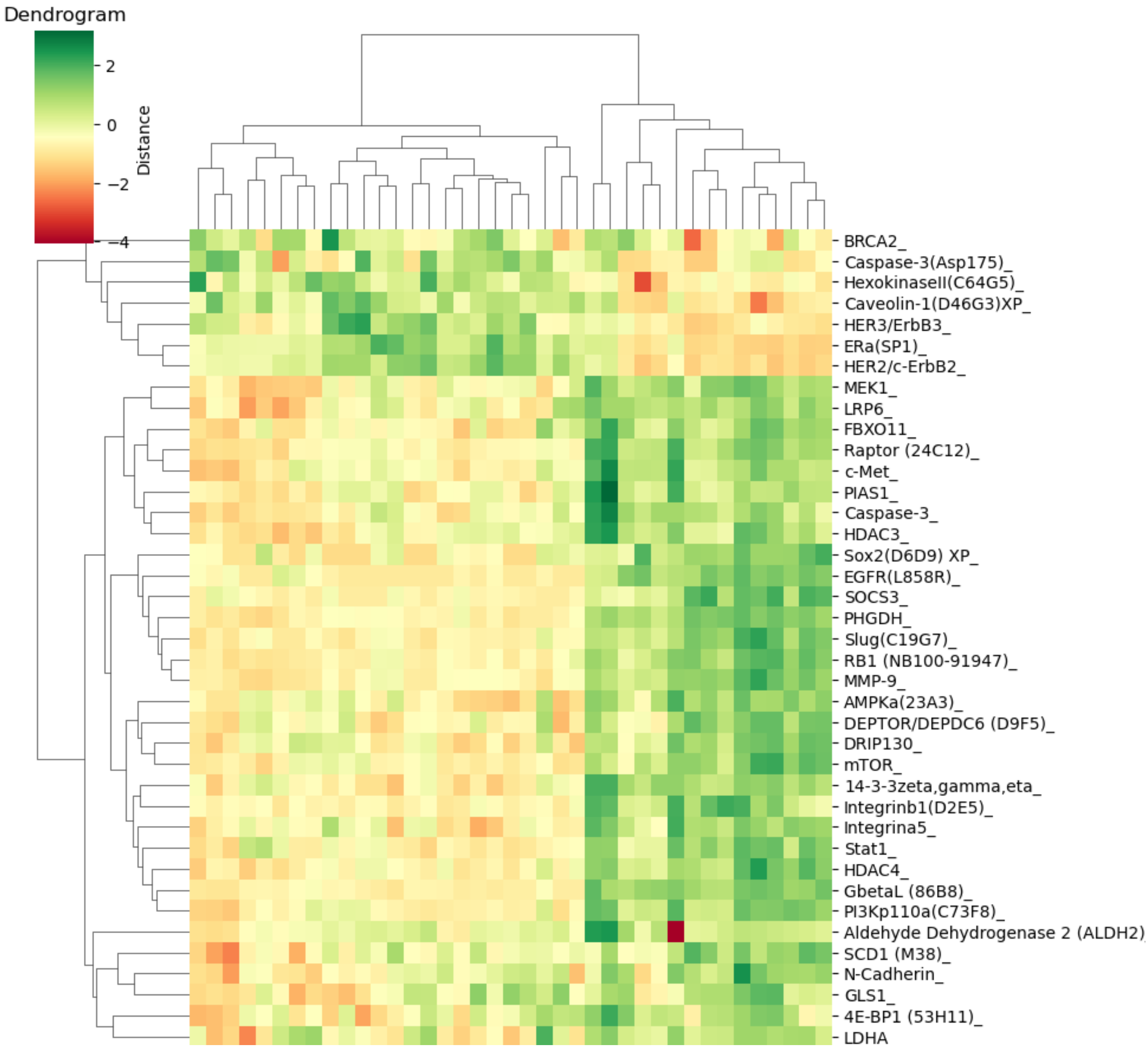


**Figure 1—figure supplement 1**

### Figure 2 suppl 1

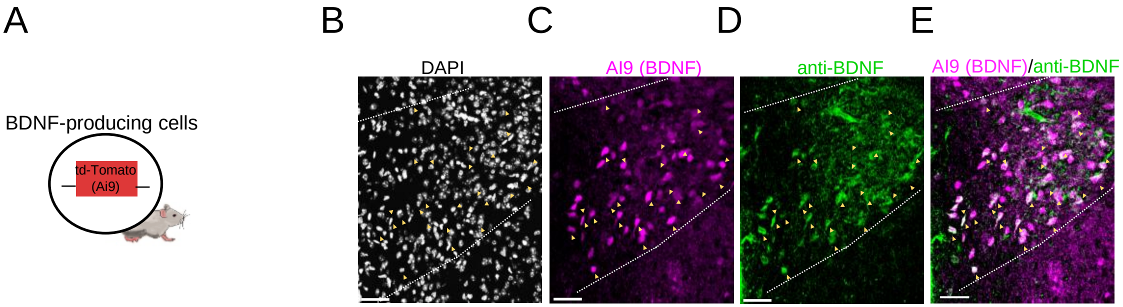


**Figure 2—figure supplement 1**

### Figure 2 suppl 2

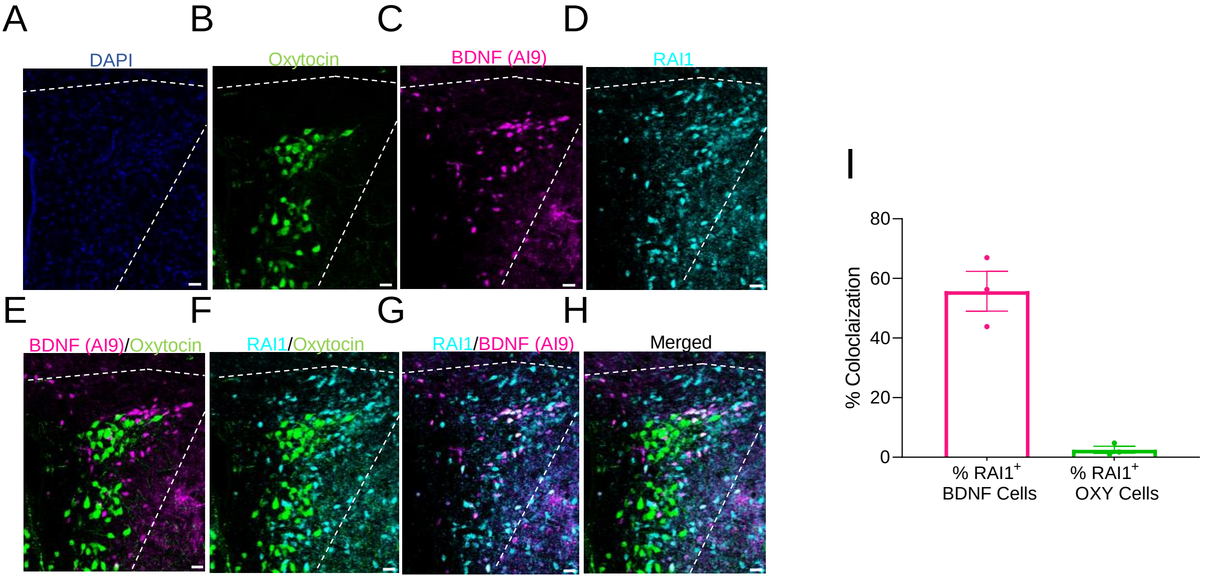


**Figure 2—figure supplement 2**

### Figure 2 suppl 3

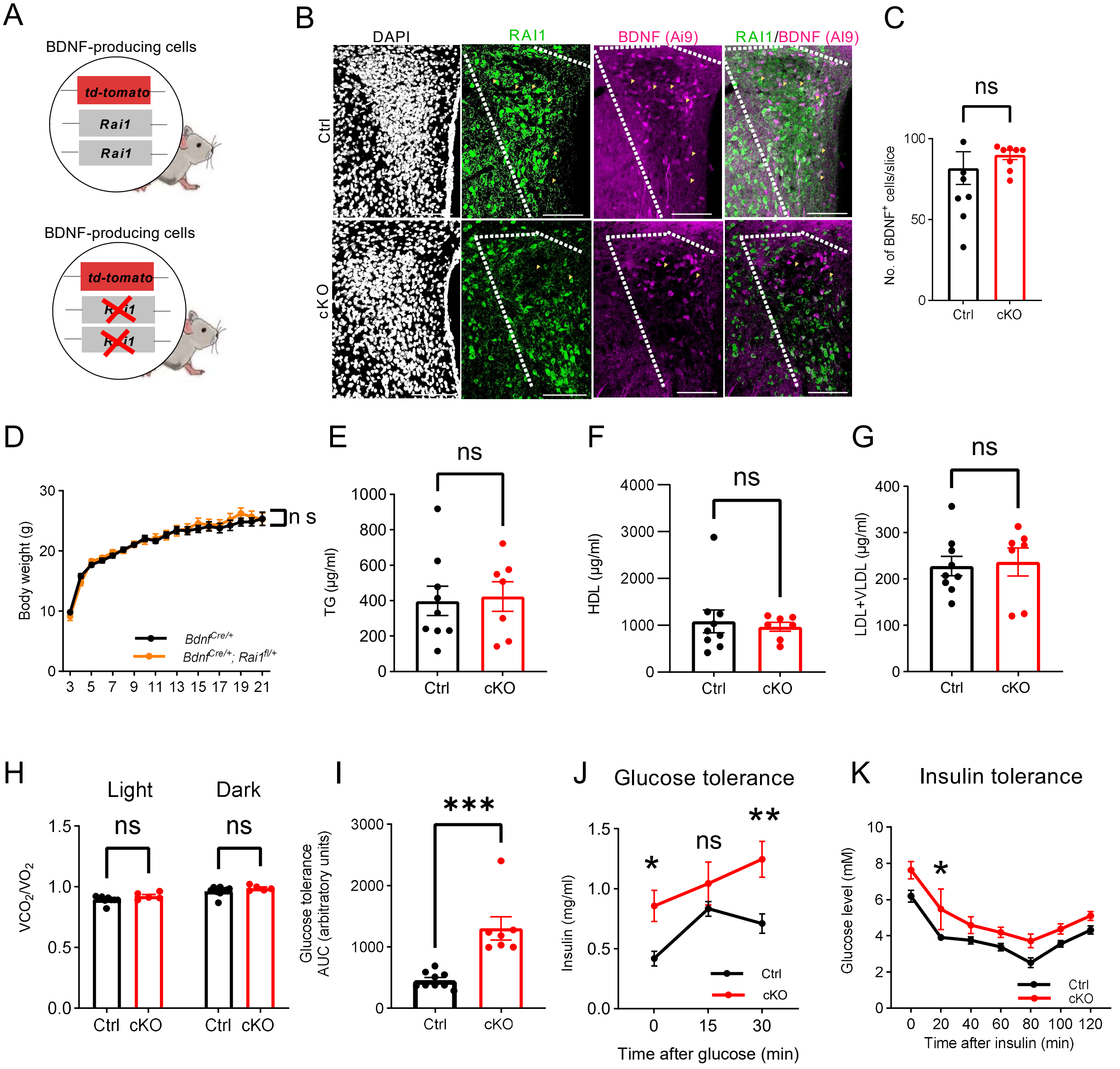


**Fig 2-figure supplement 3**

### Figure 2 suppl 4

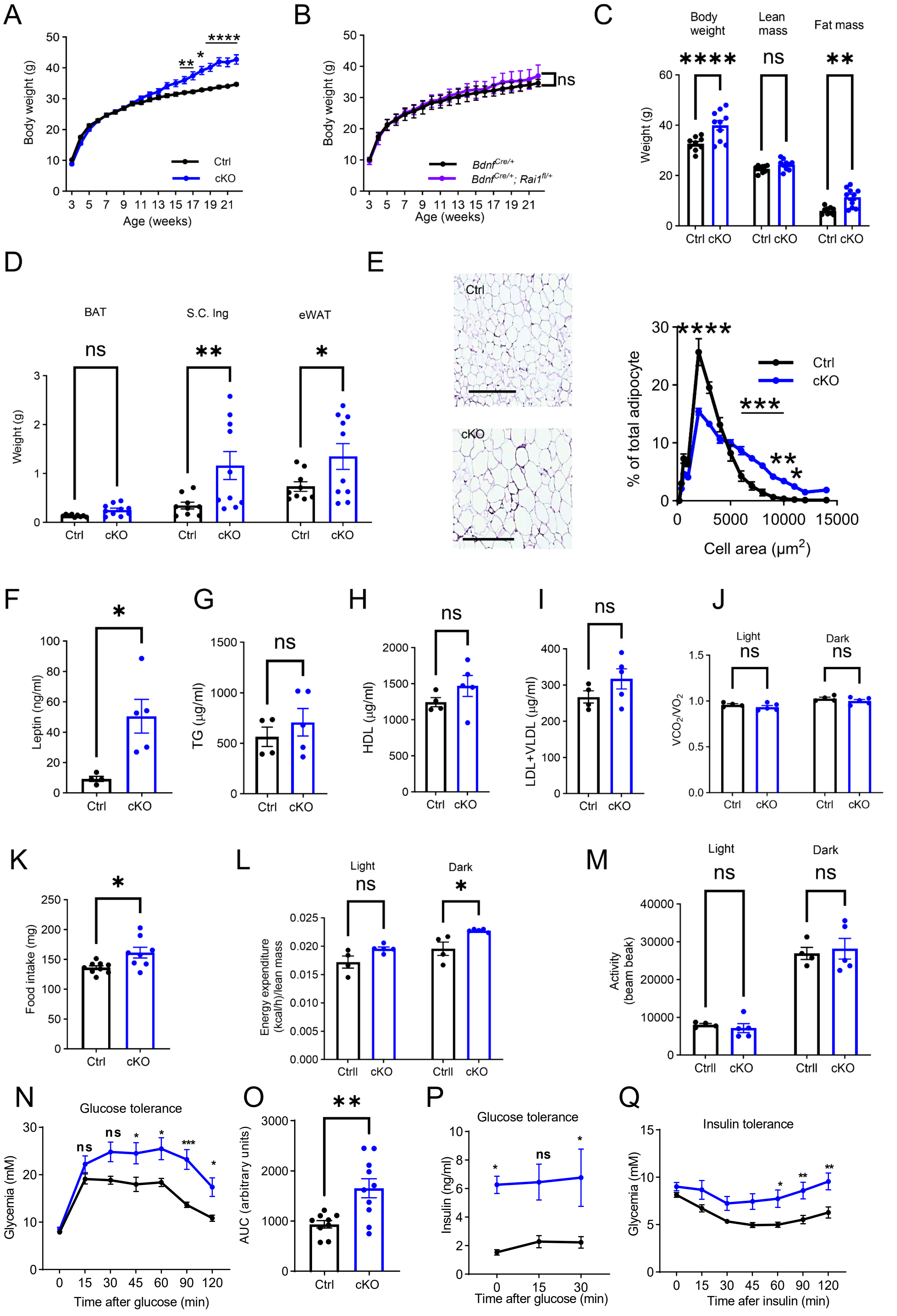


**Fig 2-figure supplement 4**

### Figure 3 suppl 1

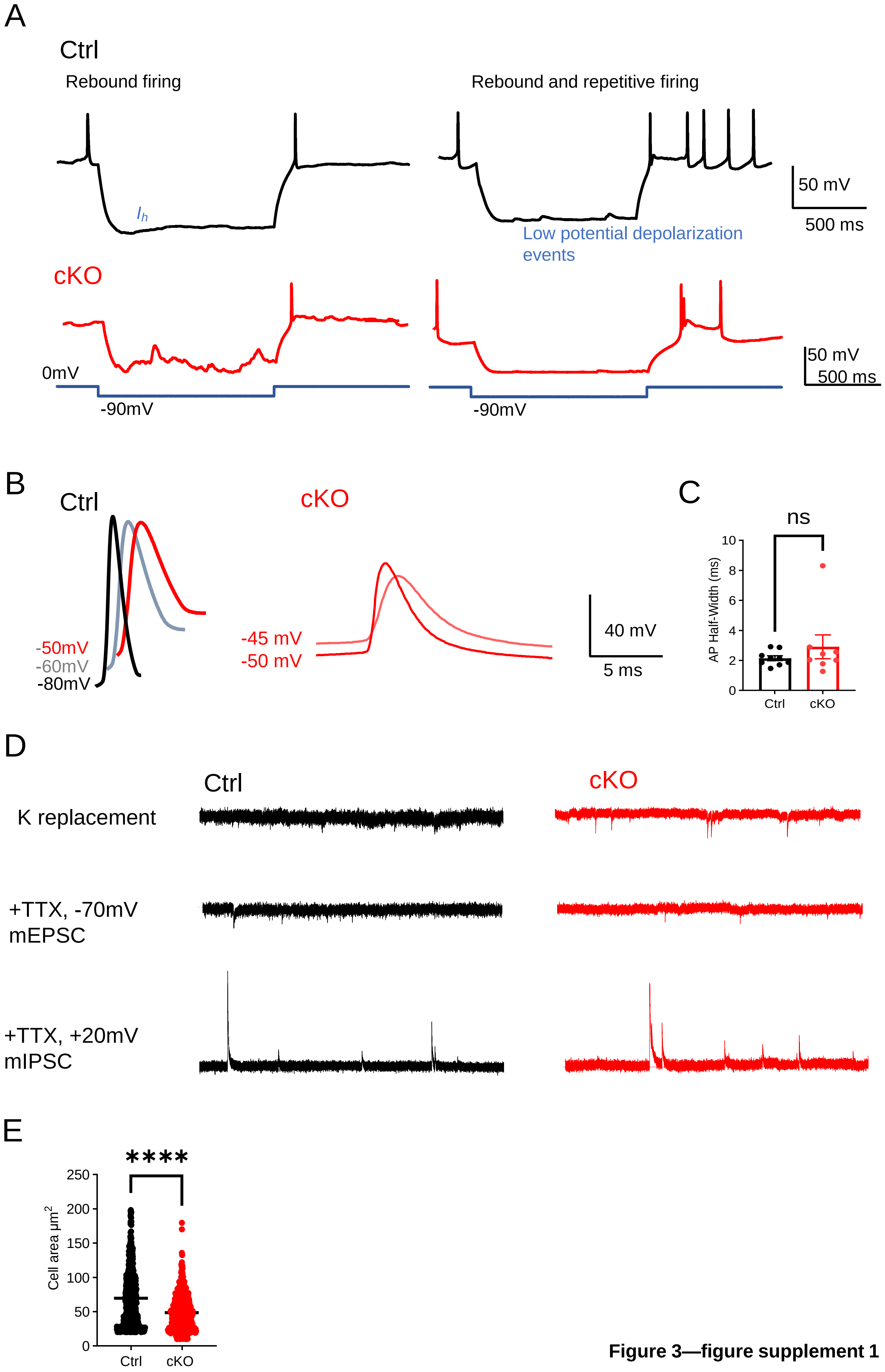


**Figure 3—figure supplement 1**

### Figure 4 suppl 1

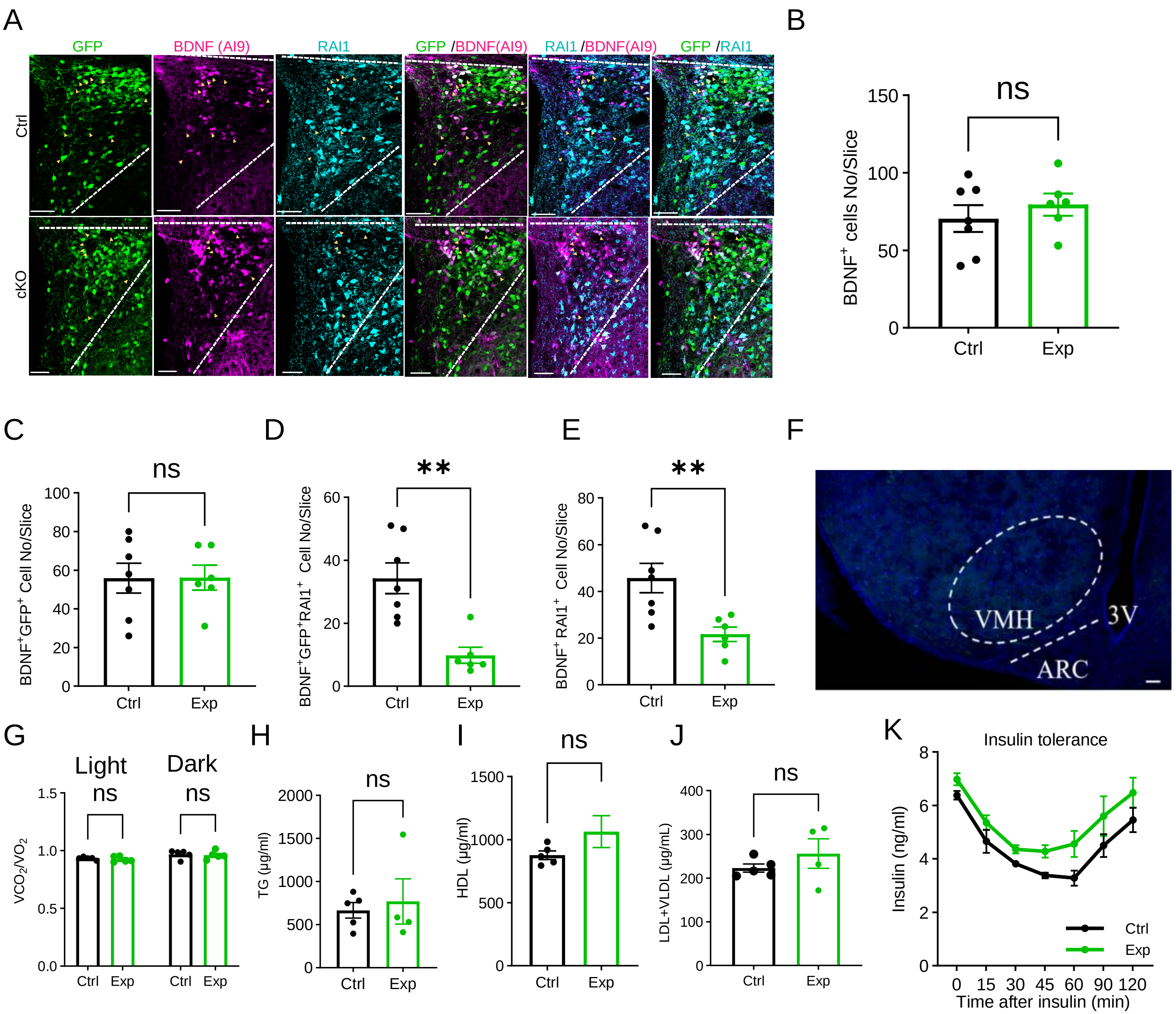


**Figure 4—figure supplement 1**

### Figure 5 suppl 1

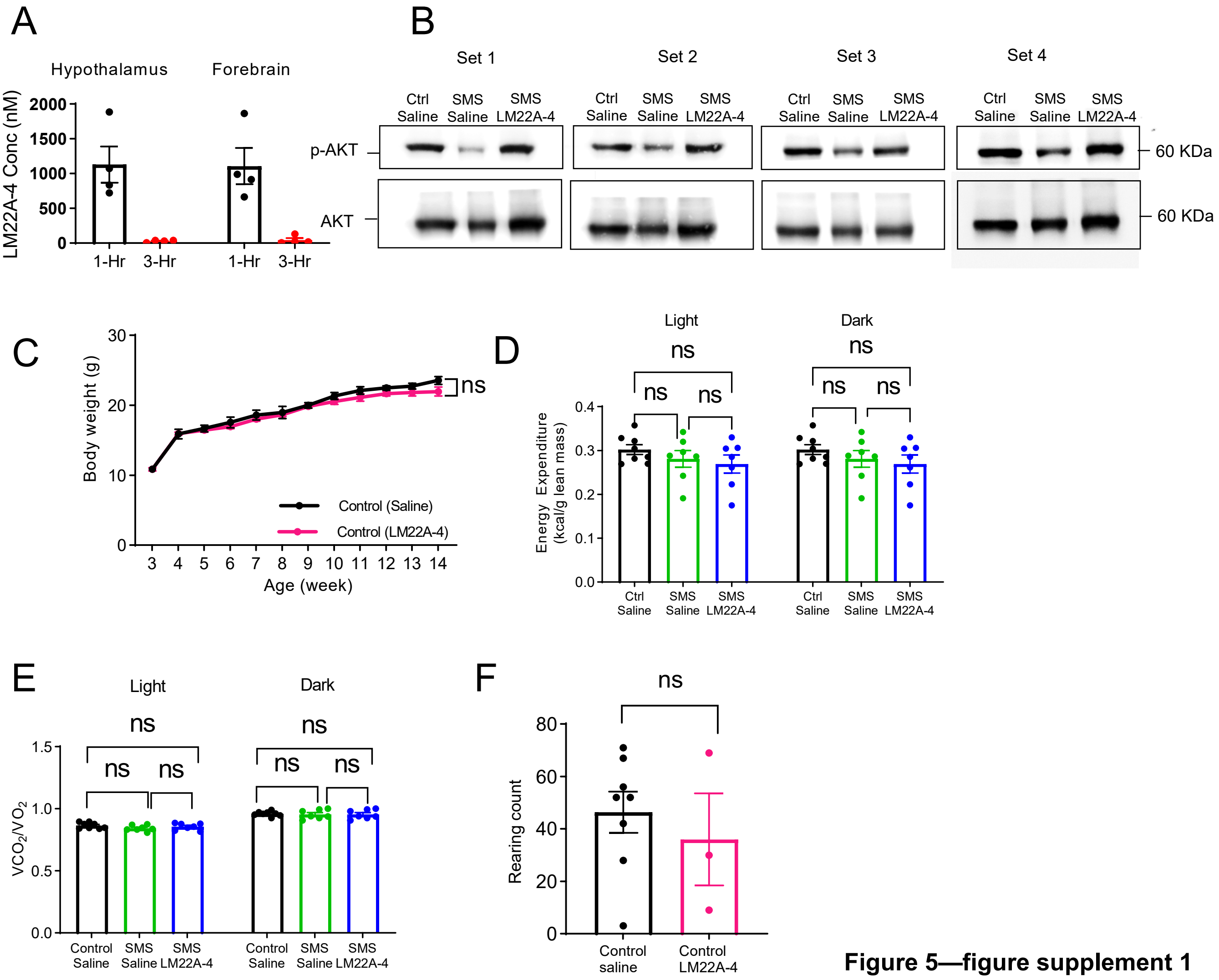


**Figure 5—figure supplement 1**

### Figure 5 suppl 2

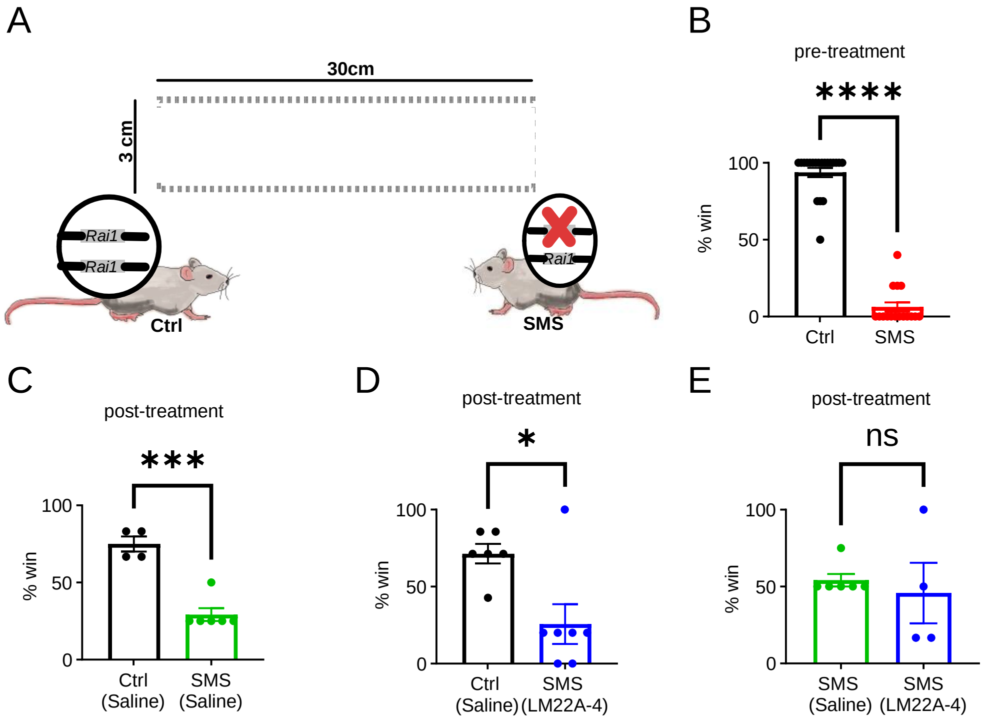


**Figure 5—figure supplement 2**
