## Supplementary material for "Smith-Magenis syndrome protein RAI1 regulates body weight homeostasis through hypothalamic BDNF-producing neurons and neurotrophin downstream signalling": western blot source data

Uncropped western blots for staining against antibodies for P-AKT & AKT. The membrane was stripped to stain for AKT after staining for p-AKT. Before staining, the membrane was cut at band 72KDa and below 53 KDa. The yellow box represents the cropped region for images shown in this manuscript.

**Set 2: Figure 5—figure supplement 1**

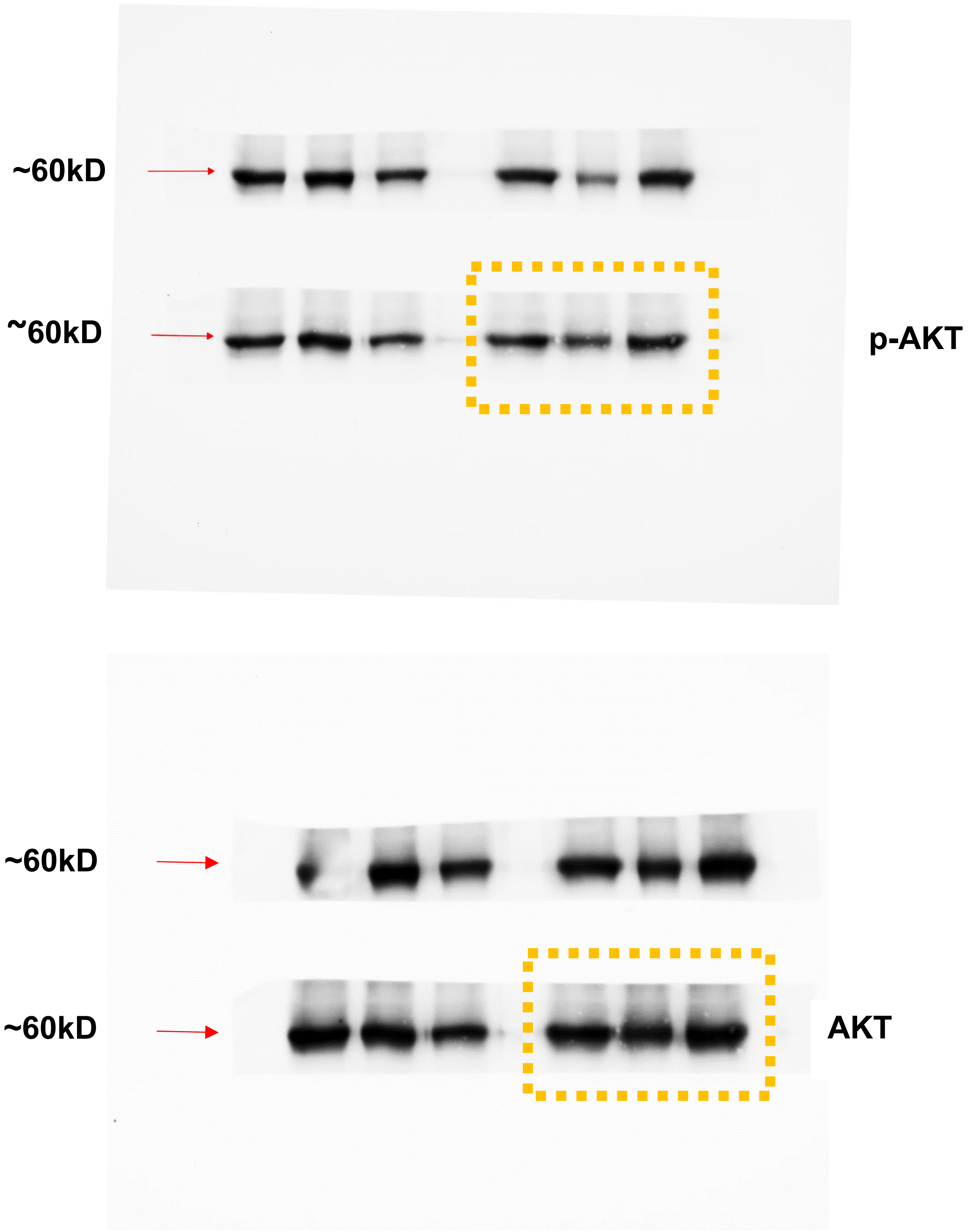

Set 4: Figure 5—figure supplement 1

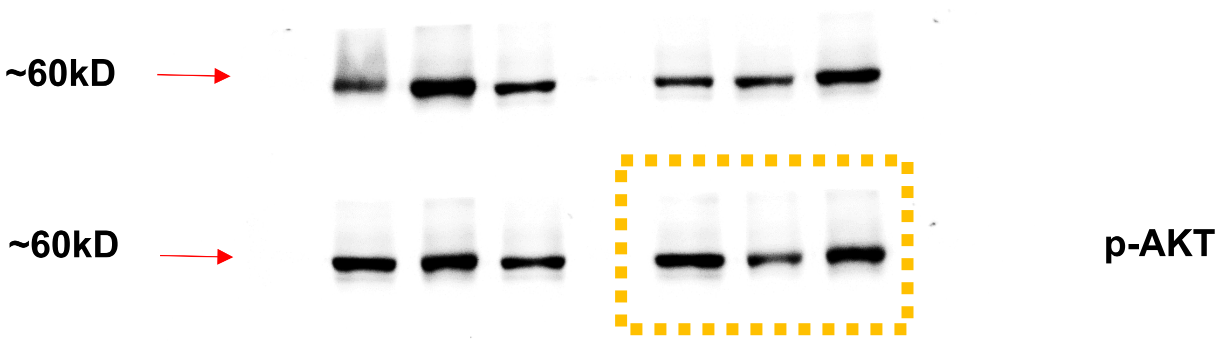


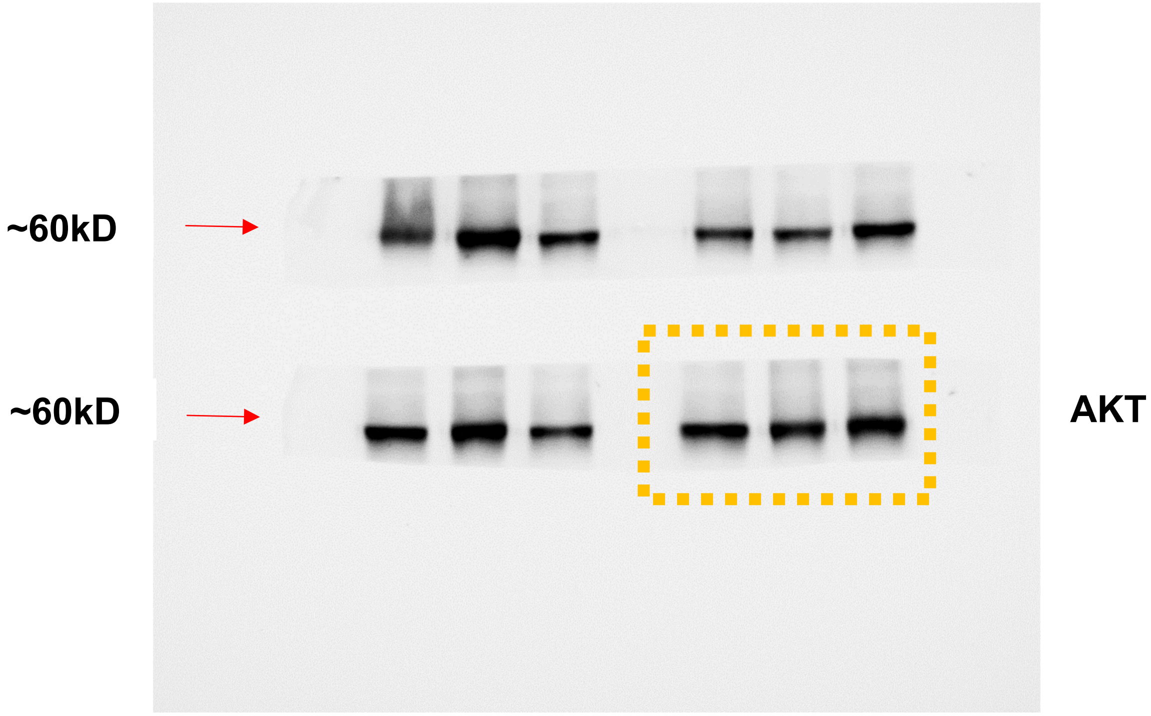

### Figure 5A, same as Set 3 in Figure 5—figure supplement 1

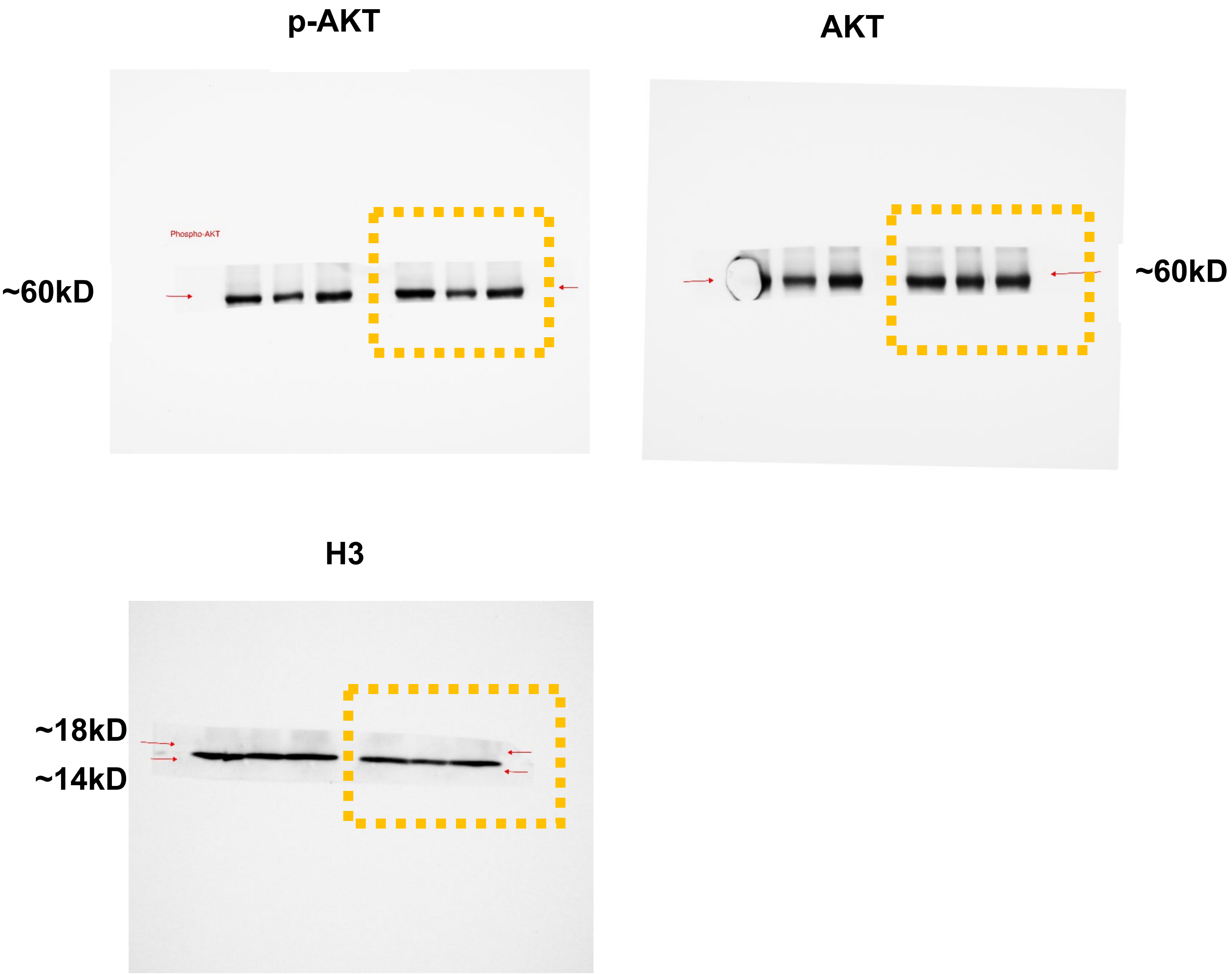


### Set 1: Figure 5—figure supplement 1

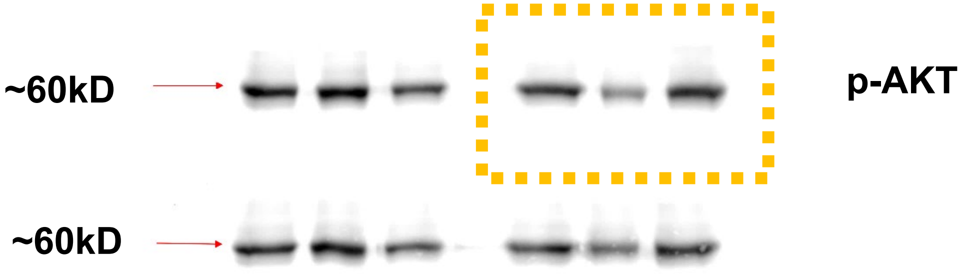

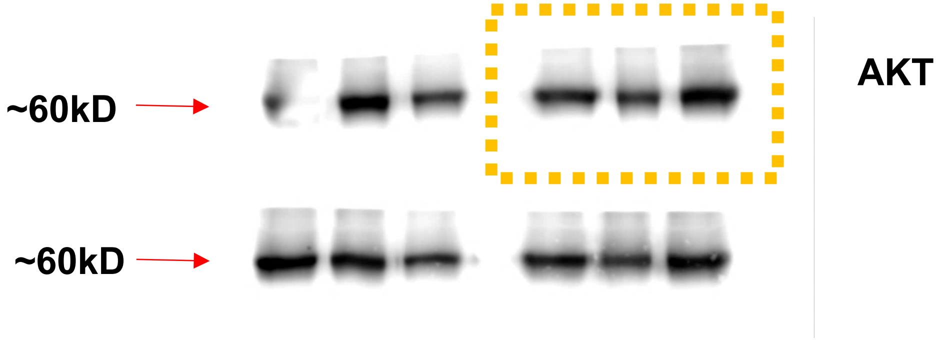
~60kD
~60kD
AKT
